# Functional impairment of the Lateral Geniculate Nucleus in Multiple Sclerosis

**DOI:** 10.1101/2022.09.12.506825

**Authors:** Neda Sardaripour, Mehrdad Asadi, Hamid Abrishami Moghaddam, Ali Khadem, Reza Rajimehr

## Abstract

Visual impairment is one of the early symptoms of Multiple sclerosis (MS) disease. The objective of this study is evaluating function of Lateral geniculate nucleus, which bridges visual information from retina to other higher order visual processing areas. We collected BOLD fMRI data from 19 MS and 19 control subjects by employing selective visual stimulation tasks to provoke the whole LGN, Magnocellular, Parvocellular, and Koniocellular pathways as part of LGN multilayer structure. Through statistical analysis, we observed a significant reduction (p<0.05) of the average BOLD signal from the whole LGN structure in MS group. Further investigations showed a significant reduction of BOLD signal (p<0.05) in response to Magno and Parvo stimuli compared to healthy controls that suggested a selective functional impairment emerging in primary visual pathways in MS. In summary, we showed functional abnormalities in LGN structure and its M and P subdivisions based on functional MRI.

## 1. Introduction

Multiple Sclerosis (MS) is characterized by self-attacking of autoimmune system to the neurons in the central and peripheral nervous system for unknown reasons, resulting in inflammation and demyelination of fiber tracts. The MS causes severe structural and functional impairments of neurons in different parts of the body and brain decreasing the transmission rate between neurons, including the optic nerve. Notably, abnormalities in visual acuity, color vision, and temporary vision loss in one eye are considered as one of the early symptoms of MS. These symptoms are the results of inflammation of optic nerves called optic neuritis (ON) (Kale, 2016).

Some previous studies observed long term structural damage and decrease of integrity because of lesion formation along the visual pathways from the retina to brain cortex (Kolappan et al., 2009, Pawlitzki et al., 2020). Atrophy of retinal layers in MS patients with (Talman et al., 2010) or without (Barnett et al., 2017; Sharma et al., 2021) ON history, indicates the spread of neuronal damage and lesion formation in both directions, *anterograde* (i.e.; toward the visual cortex), and r*etrograde* (i.e.; toward the retina), known as trans-synaptic degeneration (TSD) (Tian et al., 2018). TSD in visual pathways has been shown as one of the reasons for volume reduction of subcortical brain structures such as thalamus and its specific nuclei like lateral geniculate nucleus (LGN), (Planche et al., 2020 Papadopoulou et al., 2019), causing sustained vision disability in early MS patients (Zivadinov et al., 2013).

Using functional MRI, an extensive body of research has reported less activation in the early visual areas of the MS patients, with or without ON history, compared to healthy controls, which probably depicts reduced inputs due to structural degeneration of neurons (Faro et al., 2002; Gareau et al., 1999; Langkilde et al., 2002; Rombouts et al., 1998; Russ et al., 2002; Toosy et al., 2002; Werring et al., 2000). Notably, in severe MS patients, the brain recruits higher-order cortical areas with extra activation to maintain normal vision, known as cortical plasticity (Rocca & Filippi, 2007; Toosy et al., 2002; Werring et al., 2000). In addition to cortex, an early neuronal reorganization known as plasticity has been observed in the LGNs during recovery phase (<50 days) (Korsholm et al., 2007, Faro et al., 2002).

LGN is a fundamental thalamic center in the transmission of compressed visual information from retina to cortical areas (Briggs & Usrey, 2008; Sherman & Guillery, 2002). Therefore, focusing on LGN’s structural and functional organizations affords a useful insight into visual processing deficits in MS disease. Based on the previous studies, magnetic resonance imaging (MRI) modalities can reliably and robustly detect small subcortical nucleus like LGN in human visual system (Anderson et al., 2009; Chen et al., 1999; Chen & Zhu, 2001; Haynes et al., 2005; Kastner et al., 2004; O’Connor et al., 2002). Specifically, functional and structural MRI also enable investigating subdivisions of LGN (i.e. magno and parvo) in human brain (Denison et al., 2014; D’Souza et al., 2011; Mullen et al., 2008, 2010; Schneider et al., 2004; Zhang et al., 2015 Müller-Axt et al., 2021).

There are three parallel visual processing streams: Magnocellular (M), Parvocellular (P), and Koniocellular (K). Based on color opponent process theory, color perception stems from the interaction between three parallel visual processing streams: *red-green* (R/G), *blue-yellow* (B/Y), and *black-white* (B/W). M, P, and K pathways convey the color opponent signals from retina to higher order processing centers (Dacey, 1999; Martin et al., 1997; Mullen & Sankeralli, 1999). LGN as a thalamic multilayer structure, composed of M, P, and K layers, provides good opportunity to study these three visual pathways. These layers differ in cyto- and myelo-architecture, and in excitement properties like spatio-temporal frequencies, luminance, and chronic preferences. Contribution of LGN and its different layers in visual system has been studied in-depth in healthy brains, initiated assessing the LGN layers deficits in brain disorders which affect the visual perception, like dyslexia (Ahmadi et al., 2015), glaucoma (Zhang et al., 2016), schizophrenia and mood disorders (Dorph-Petersen et al., 2009). In this regard, much is known about the structural changes of primary visual pathways in MS. To the best of our knowledge, discovering functional abnormality in LGN subdivisions based on functional MRI remains unaddressed in MS patients.

The aim of this study is to explore malfunction of LGN region in MS and selective abnormalities emerging in M, P, and K visual pathways located in the LGN multilayer structure. We measure the function of M, P, and K layers in LGN by provoking and receiving the BOLD fMRI signals using selective stimulation patterns, i.e. LGN localizer checkerboard and specific Magno, Parvo, and Konio stimulation patterns. We then perform statistical analysis on the preprocessed fMRI data to compare the LGN function of MS patients with those of healthy controls.

## 2. Materials and Methods

### 2.1. Background: The Lateral Geniculate Nucleus and visual pathways

In the hierarchical structure of the visual system, compressed retinal information are transferred through optic tracts to the optic chiasm, and synapsed in the LGN, located in thalamus. Fig 1-A illustrates the multilayer structure of LGN composed of M, P, and K layers. M and P are topographically located in inferior and superior organized layers, respectively. K is in the form of thinner K layers spreading between M and P layers. The left and right LGNs modulate the visual information to the upper-level processing cortical regions through optic radiation (OR) bundles formed by axonal processes of M, P, K cells. So, each LGN is characterized as a central node linking the anterior and posterior visual pathways. In addition to the relay function of LGN, there is a bilateral interaction between thalamus and cortex (Schmielau & Singer, 1977) in which, LGN receives feedback from other processing areas of the brain like V1 and Superior Colliculus (SC) to modulate inputs to the higher visual processing centers (Kastner et al., 2006). Based on color opponent process theory, the color perception stems from the interaction between three parallel visual processing streams: *black-white* (B/W), *red-green* (R/G), and *blue-yellow* (B/Y). These pathways convey the color opponent signals from retina to higher order processing centers (Dacey, 1999; Martin et al., 1997; Mullen & Sankeralli, 1999).

**Figure 1.**
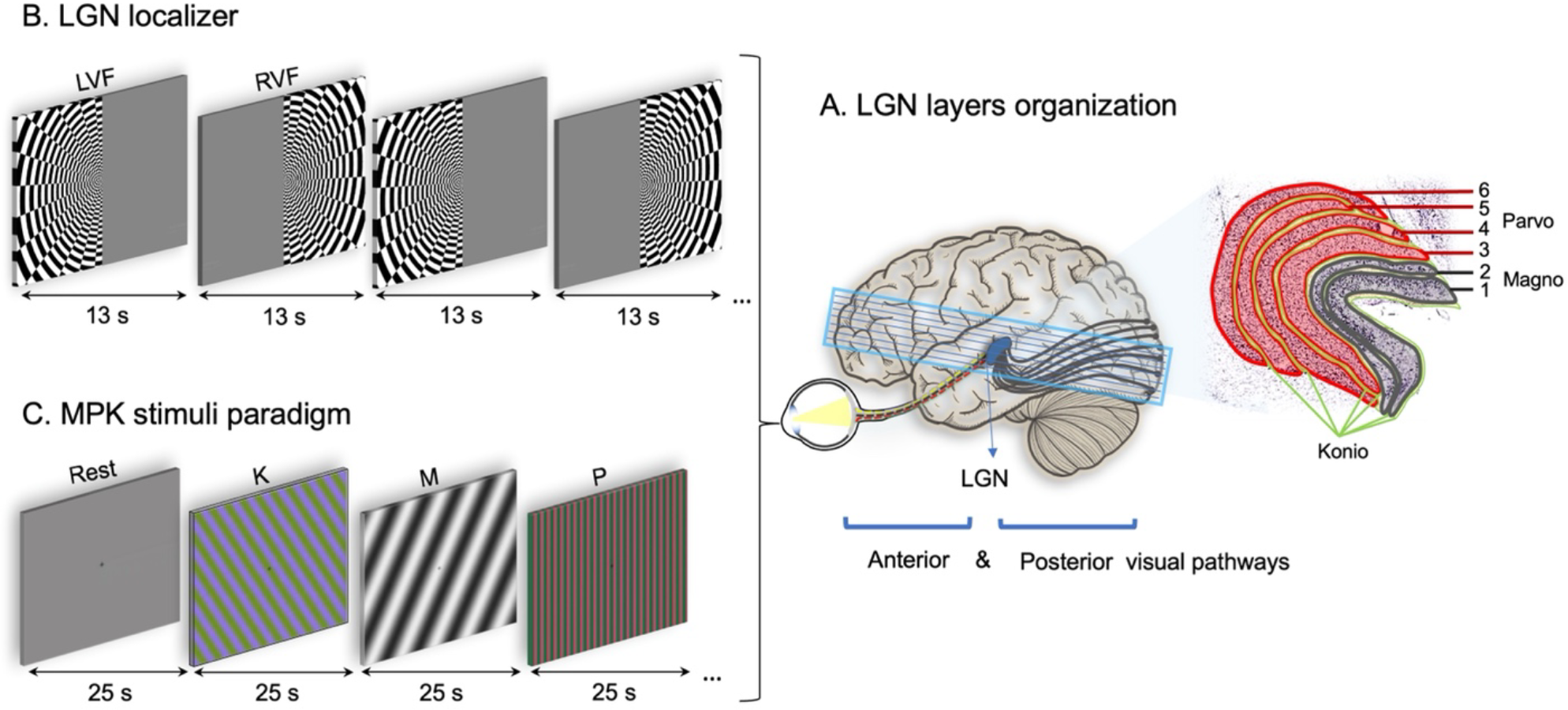
Visual pathway stimulation block diagram. Neural stream of primary visual pathways from retina to cortex and the multilayer anatomy of LGN region. Noted, non-whole brain fMRI volumes with restricted ROI were acquired. The Blue color border denotes the slice prescription during data acquisition from sagittal view (A). The LGN localizer pattern in form of left visual field (LVF) and right visual field (RVF) blocks (B). Magno, Parvo, and Konio stimulation blocks, noted by M, P, and K (C).

### 2.2. Visual stimulus characteristics

In this paper, we propose two simulation paradigms. First, to localize the whole LGN region, we generate a hemifield black/white flickering checkerboard pattern alternating between left and right visual fields (LVF and RVF) every 13 seconds, as shown in Fig 1-B. The *LGN localizer* has a contrast of 100% reversing with the frequency of 4Hz. The square size of checkerboard pattern increases exponentially by increasing distance from the center.

Second, to stimulate the LGN sublayers, we generate an *MPK stimuli paradigm* consisting of four blocks each lasting 25 seconds, namely *M* (for evaluating non-color vision), *P* and *K* (for evaluating color vision), and *Rest* (resting state gray block), as shown in Fig 1-C. Each block is rotated randomly in six angles (0, 30, 60, 90, 120, 150) every 3 seconds. The order of M, P, and K blocks changed randomly between every two consecutive Rest blocks. Moreover, the subjects are asked to focus on the “+” sign displayed at the center of each block.

The M and P blocks and LGN localizer checkerboard were inspired from (Denison et al., 2014). We added K blocks based on color and spatiotemporal characteristics of Konio cells. We anticipate that this K block can distinguish the function of K pathways from M, and P, enabaling the evaluation of K layer dysfunction in MS patients (Yoonessi, 2011).

M pathways are stimulated by an achromatic, high luminance contrast, low spatial (0.5 cpd) and high temporal frequency (15 Hz) stimulus. P pathways are provoked by isoluminant chromatic (R/G), high color contrast, high spatial (2 cpd) and low temporal frequency (5 Hz) stimulus. The spatiotemporal characteristics for provoking K pathways are between these two, with spatial frequency of 1 cpd, temporal frequency of 10 Hz, high color contrast and isoluminant chromatic (yellowish green-bluish purple) pattern.

### 2.3. Stimulation procedure

We generate the stimuli by Psychopy software (Version 3.0.7), using Python. A projector (frame rate=60Hz) located outside the scanner projects the stimulation in full screen mode on a white screen placed at the end of the scanner bore behind the subject’s head. The stimulation is reflected via a mirror mounted over the subject’s eyes. For correct color representation, the room light was turned off during the fMRI session.

For each subject, we acquired fMRI data in two functional runs: 1) the *LGN localizer* run containing 11 periods of left visual field (LVF) and right visual fields (RVF), and 2) the *MPK stimuli paradigm* run containing M, P, K, and Rest blocks (25 blocks each). We collected 119 and 260 fMRI volumes during LGN localizer and MPK stimuli paradigm, respectively. In total, functional data collection took ~15 min for each subject.

### 2.4. Data collection

#### 2.4.1. Subjects

We collected a task-related fMRI dataset comprising 21 MS patients (8 males, age 32.47 ± 5.48) and 21 healthy controls (9 males, age 32.13 ± 4.81) from Iran National Brain Mapping Laboratory (NBML) and Kasra Hospital. Two patients and two healthy control were excluded after checking data quality during preprocessing steps.

The average MS duration among the patients was 10 years. Nine patients had no ON history (ON−) while the rest of them had previous ON (ON+) as a part of relapsing remitting MS. This experiment was completely approved by the Iran University of Medical Science ethical committee (IR.IUMS.REC.1395.9301984). A consent was acquired from each participant before the experiment. Before starting data acquisition, a physician examined the subjects for the following inclusion criteria:

1. Absence of visual problems caused by MS influencing the visual pathway’s normal function, such as blindness, low vision, and blurred vision (previous ON should occur at least 3 months before the data collection),
2. Absence of eye or brain surgery,
3. Absence of any metal object such as platinum, artificial heart valve, pacemaker, brain shunt tube, metal chips or eyes/ears implants,
4. Normal cognitive function for staying concentrated during visual tasks.

#### 2.4.2. MRI imaging protocol

Task-related fMRI from 16 MS patients and 16 control individuals were acquired via the 3 Tesla SIEMENS MAGNETOM Prisma scanner in NBML, while the rest were scanned via the 1.5 Tesla SIEMENS MAGNETOM Avanto syngo MR B19 scanner in Kasra hospital.

Before applying visual stimulation, we recorded field-map, and high-resolution T1-weighted structural MRI for preprocessing purposes. Moreover, a single whole-brain volume fMRI data was also acquired to improve the registration. During both MPK and LGN localizer stimuli, we acquired non-whole brain fMRI volumes with restricted ROI as shown in Fig. 1-A in blue color. The whole process of data acquisition from preparation to the end of the imaging took about 45 min for each subject. For more details about the imaging parameters are provided in the Appendix 1.

### 2.5. Data analysis

Our goal is to investigate the LGN function and its layers (M, P, and K) in MS patients using the stimulation paradigms presented in Section 2.2. To do so, we first perform the preprocessing pipeline (see Section 2.5.1) on the collected fMRI data, and then the statistical analysis (see Section 2.5.2).

We localize the functional boundaries of LGN region in both MS and heathy groups using the statistical analysis on fMRI data acquired during LGN localizer stimulus (Fig 1-B). The resulted activation map from healthy group will be used as the binary mask to apply for the LGN ROI analysis. Then we perform the statistical analysis on fMRI data acquired during MPK stimuli paradigm to evaluate LGN layers function in MS patients.

We used FreeSurfer and MATLAB R2017a software suites, and to visualize the volume-based results we used TkMedit package. The following subsections provide more details about the data analysis.

#### 2.5.1. Preprocessing

As the data preparation steps, cortical reconstruction and volumetric segmentation processes were applied on 3D structural data in FreeSurfer. They include motion correction, non-brain tissue removal, automated Talairach transformation, volumetric white matter (WM) and gray matter (GM) segmentation, intensity normalization, GM and WM boundaries tessellation, topology correction, and surface deformation procedures. Next, we used FS-FAST in FreeSurfer for preprocessing steps. Also, we considered other software suite algorithms to improve some preprocessing steps. The preprocessing pipeline includes Motion Correction (using AFNI), Registration (Choosing the best output of FSL, SPM and FreeSurfer algorithms for each subject), slice timing correction, spatial smoothing (FWHM= 3 mm, to reduce inter-subject variability), brain masking (using FSL’s BET), and normalization. It is noteworthy that to improve the quality of structural on functional co-registration, we used a single whole-brain volume fMRI data as an intermediate stage. After completing the preprocessing, we started the statistical analysis.

#### 2.5.2. Statistical analysis

We performed volume-based statistical analysis to extract significantly activated voxels during the stimulation. In particular, we first constructed a *timing matrix* containing labels, start time, and duration of LVF and RVF blocks as presented in LGN localizer paradigm. Similarly, a timing matrix for M, P, and K blocks in MPK stimuli paradigm was constructed. Next, we defined several *contrast matrices* for both paradigms including M-contrast (Magno vs. Rest), P-contrast (Parvo vs. Rest), K-contrast (Konio vs. Rest), PK-contrast (Parvo vs. Konio), and LR-contrast (LVF vs. RVF).

Given the motion regressors estimated from preprocessing step, timing, and contrast matrices, we applied generalized linear model (GLM) to best fit the preprocessed fMRI timeseries to the expected voxel timeseries which was obtained based on hemodynamic response function (HRF) and timing matrices. After estimating beta-values for each condition, and given the defined contrasts, we obtained the statistically significant (p-value<0.05) activation maps for all subjects. We applied the binary mask obtained from LGN localizer stimuli to calculate the average amplitude of BOLD signal for each subject. Next, we defined a contrast matrix for entire MS and healthy groups and performed weighted random effect (WRE) model over entire population to identify clusters of significantly activated voxels (p-value<0.05) for each group. Finally, a between-group comparison identified the cluster of voxels which were showing a statistically significant difference (p<0.05) between two groups. We performed a cluster-wise correction for multiple comparison by running a permutation simulation and get the cluster-wise corrected p-values (CWP).

## 3. Results

### 3.1. Localizing the LGN region

To assess the whole LGN region function, we presented the checkerboard localizer to subjects and followed the processing pipeline as explained in Section 2.5.

The group functional statistic map was identified by contrasting between fMRI responses to the M and P stimuli (paired t-test across all subjects). Fig 2-A shows the functional activation map of the whole LGN region in both healthy and MS groups during LVF and RVF checkerboard stimulation blocks. The boundaries of LGN depicted in green color are identified based on the physiologically expected anatomical location of LGN on high-resolution T1-w images (Zhang et al., 2015b). The statistically significant clusters (p<0.05) were identified by contrasting between fMRI responses to the RVF and LVF stimulation blocks. The resulted statistical maps from the LGN localizer have overlapped with the depicted ROI. Moreover, consistent with the visual system anatomy, the LVF and RVF checkerboard stimulus have activated right and left LGN as showed by blue and red color, respectively.

**Figure 2.**
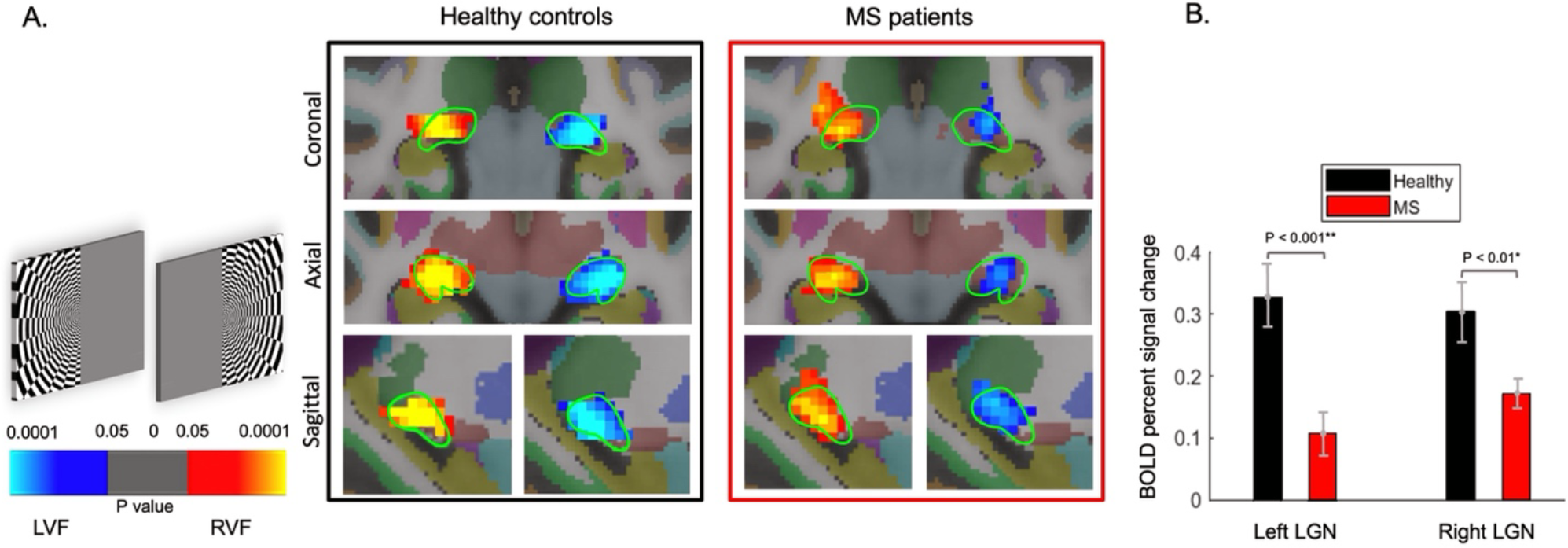
LGN Localizer activation map. (A) The cluster of significant active voxels (p<0.05) overlapped with the physiologically expected locations of LGN depicted by green lines (the LGN boundaries in right column match exactly with the left one), which illustrates the location and shape of the human LGNs were defined from T1 structural images. The left and right columns represent the LGN functional map of healthy control and MS groups, respectively. The MNI coordinates of the centers of mass of the significant clusters after multiple correction (CWP<0.0001) were: right control (24.00, −25.00, −7.00), left control (−26.00, −25.00, −7.00), right MS (20.00, −29.00, −3.00), and left MS (non-significant cluster after correction). (B) Bar plot shows the difference of healthy and MS groups, in form of BOLD signal percent change in left and right LGN, separately. Error bars in this figure indicate standard errors of the mean.

Fig 2-B shows the BOLD percent signal changes and statistical difference of both groups in left, and right LGN separately. It is clear that LGN activation in MS group has declined in both hemispheres compared to healthy controls.

### 3.2. The evaluation of LGN sublayers function in response to MPK stimulation

To evaluate the function of three visual pathways located as the sublayers of LGN region, we first presented the *MPK stimuli* consisting of luminant achromatic (M), and isoluminant chromatic (P and K) blocks to subjects, as shown in Fig 1-A. After completing the preprocessing steps, and statistical analysis on the LGN sublayers, the significant active voxels were extracted in response to MPK stimuli paradigm. Fig 3 indicates the average of BOLD signal in response to each of M, P, and K stimuli in both groups. The Magno and Parvo pathways show significant lower BOLD signal in MS group. Although the BOLD average signal of Konio layers in both left and right LGNs is lower than healthy controls, the response reduction is not statistically significant (p>0.05). We figure out the functional pattern in response to K stimulus is not statistically nor topologically different than the P functional map as the P-K contrast resulted no significant active voxel.

**Figure 3.**
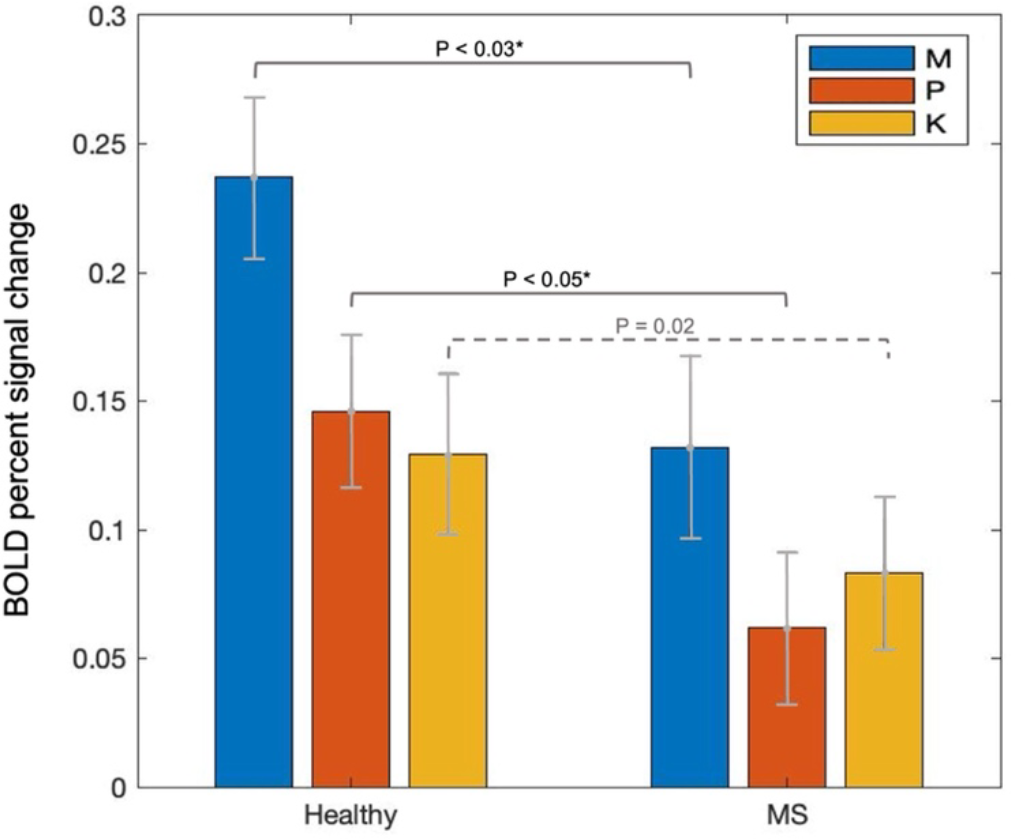
BOLD percent signal change acquired from the LGN region in both healthy and MS groups, in response to M, P, and K stimuli. The average BOLD signal in MS groups show significantly lower value compared to healthy group (p<0.05), indicating the functional impairment of the Magno and Parvo layers due to MS disease. Error bars indicate standard errors of the mean.

Given the statistical results in this study, our K stimulus failed to provoke thin and sparse K layers distinctly. P and K stimulation blocks have not showed distinguished performance from each other in provoking the LGN color processing subdivisions, as the PK-contrast showed no significant active voxel even in healthy group. So, we can state that both of P and K stimuli majorly provoke the visual pathways which convey the color information in LGN. We will further discuss our findings in the next section.

## 4. Discussion

During vision process, the information of rod and cone photoreceptors is compressed in retina in form of three distinct visual pathways and then is transmitted to the upper-level processing centers like LGN. LGN layers are the location of parallel visual information processing streams composed of three types of anatomically distinct cells – M, P, and K. M and P cells are organized in distinct spatial positions, while K cells spreads sparsely in form of thin layers between M and P layers. Although fMRI can provide a map of the neural activities in LGN organization, there is no fMRI study on malfunctions of these pathways in MS patients yet.

Based on previous studies, investigating subcortical pathways gives us insight toward human visual perception and cognition (Zhang et al., 2015b), hence it may also provide an effective way to detect and address upcoming issues and abnormalities emerging in certain neurodegenerative diseases. Notably, Sun et al. (Sun et al., 2019) proposed a hypothesis that dysfunctionality in subcortical visual pathways may be a new approach to detect visual impairment in glaucoma in early stage. On the other hand, due to the growing number of MS patients all over the world in the recent decade, MS disease has been extensively studied. It has been reported that MS patients experience cognitive disabilities and vision problems. So, it’s meaningful to evaluate LGN region and its sublayers as an important subcortical nucleus in visual system.

We employed specific visual stimulation patterns with different spatial and temporal frequencies to stimulate the whole LGN region and each pathway separately. We employed two visual stimulation paradigms, i.e., the checkerboard pattern for localizing whole LGN and MPK stimulation paradigm to provoke color and non-color sensitive pathways. Then we used an appropriate fMRI imaging protocol to get high spatial and temporal resolutions and collected fMRI data to noninvasively map the whole LGN and provoke its M, P, and K subdivisions which are part of visual subcortical pathways. We found the corresponding functional maps of significant active voxels and discussed our findings as follows:

### The significant reduction of BOLD response in LGN in MS vs. control group

Our results showed reduction of average BOLD signal and reduced number of active voxels in MS patients vs. healthy controls, suggesting functional impairment of the whole LGN in MS disease. Previous structural MRI studies observed shrinkage of the entire thalamus and its specific nuclei like LGN in MS patients (Planche et al., 2020 Papadopoulou et al., 2019), which has an association with spread of neural degeneration through primary visual pathways (trans-synaptic degeneration (TSD)). Reducing functional response in this experiment might be a result of shrinkage of LGN structure due to MS disease, or even reduction of LGN input given the TSD through the primary visual pathways from retina to cortex.

Korsholm et al., (Korsholm et al., 2007) suggested that a very early recovery takes place in optic nerve following an episode of acute optic neuritis (ON) in less than 50 days (~2 month). Our study was focused on MS patients without visual problem and history of ON at least 3 months before fMRI data acquisition. However, our findings showed decreased LGN activation in MS patients suggested that a substantial remyelination process does not occur during recovery in MS which leads to irreversible damage to neurons. Our finding is in agreement with previous studies which reported reduced amplitude and prolonged latency in visual evoked potential (VEP) in MS (Pihl-Jensen et al., 2017). Further exploration is needed in future studies to find out the interplay between demyelination and recovering remyelination by using functional-structural data in MS.

Beside the significant reduction of BOLD signal in the whole LGN body, we aimed to assess the selective functional impairment of LGN sublayers in MS patients. So we used M (black-white), P (Green-Red) and K (Yellowish green-Bluish purple) block patterns that could be categorized into two groups of luminant achromatic (M) and isoluminant chromatic (P and K) visual stimuli. We will discuss our findings as follows:

### The significant reduction of BOLD response in M layers of LGN in MS vs. control group

We provoked M pathways using M blocks with specific characteristics like B/W high luminance contrast, low spatial (0.5 Cpd) and high temporal (15 Hz) frequencies. After performing statistical analysis, we assigned the largest cluster of significant active voxels to M layers located in the LGN ROI mask. Our results showed reduction of average BOLD signal in MS patients in response to M stimulation pattern, suggesting functional impairment of magnocellular pathways in LGN compared to healthy individuals. This finding agrees with a previous MS study using evoked potential data (Murav’eva et al., 2009) which reported that in MS patients with visual impairment at low spatial frequencies, M cells have been structurally damaged by observed lesions.

### The significant reduction of BOLD response in P/K layers of LGN in MS vs. control group

We used two grating patterns: *red-green* (R/G), and *blue-yellow* (B/Y), with specific temporal and spatial frequencies to examine the color vision in MS patients. It is notheworthy that Parvo cells are smaller in size compared to Magno cells. Our finding indicated a statistically significant reduction of BOLD signal in response to P stimulus in MS patients compared to controls. In contrast, the average BOLD signal was not significantly different from the control population in response to K stimulus.

An autopsy study proposed that smaller neurons with smaller axons in LGN may be more prone to selective atrophy in MS disease (Evangelou et al., 2001). In more details, this study assessed the anterior visual pathways of eight MS patients after death to explore MS-associated structural changes in neurons. They reported that the size distribution of Magno cells was unchanged, but the size distribution of Parvo cells has a large variation in MS group. Given the TSD process, atrophy of parvo cells leads to the axonal loss of PC pathways before synapsing into the LGN layers. This process would justify decreased activity in P layers of LGN, which is consistent with our finding in this fMRI study.

A decrease in average bold signal in the MS group is also observed in response to the K stimulus compared to the healthy group. However, the difference is not statistically meaningful. This contradicts our expectation to observe K-layers malfunction in MS patients as the Konio cells are smaller than Parvo cells and more vulnerable to atrophy in MS (Evangelou et al., 2001). Although the M, P, and K visual pathways are spatially distinct in the cortex, they are closely interrelated at the pre-cortical stage. According to previous physiological studies, the M, P, and K layers in the LGN region are provoked by a combination of retinal photoreceptor output. Moreover, the frequency sensitivity of P and K streams are similar to each other (McKeefry et al., 2001; Wesner & Brazeau, 2019). Therefore, both K and P may be provoked in response to chromatic visual stimulation. In our experiment, the spatiotemporal characteristics we chose to provoke K pathways (1 Cpd, and 10 Hz) were between M (0.5 Cpd, and 15 Hz) and P (2 Cpd, and 5 Hz). This may slightly excite M layers as well. Consequently, the activation map obtained from PK-contrast (P block vs. K block) in our experiment showed no significant active voxel in the healthy group, indicating that the K stimulation block has not distinctly provoked K layers.

Notably, Konio cells in LGN are sparsely organized in thin layers between M and P; as such, detecting the response of K layers and exploring their malfunction patterns in MS requires a stronger MRI scanner (7 Tesla). Modifying the K block toward distinguishing P and K and increasing the number of subjects in our experiment are another ongoing investigation by our team. In the following, we discuss the spatial organization of our obtained statistic functional maps from the topological perspective. According to Fig. 2, the functional activation map of whole LGN region in both healthy and MS groups which was identified by contrasting between fMRI responses to the RLV and LVF stimulation blocks has completely overlayed within the boundaries of LGN (green color lines), which were identified based on the physiologically expected anatomical location of LGN on high-resolution T1 data. Moreover, according to the visual system anatomy, the visual information in the LVF projects to the right LGN, and the visual information in the RVF projects to the left LGN. This is consistent with our LGN localizer result maps, in which the LVF has activated right LGN showed by blue color, and RVF has activated left LGN showed by red color.

Figure 4 shows the spatial pattern of M and P layers in control and MS groups. The group functional statistic map identified by M-contrast, and P-contrast of fMRI responses, is overlayed on the structural T1 image in MNI space and a topographic map of the LGN layers as a reference. The color bar at the left indicates the P-values. The yellow-red colors depict statistically significant voxels (P<0.05) in response to only M block, and the blue color shade shows the significant voxels (P<0.05) in response to only P block.

**Figure 4.**
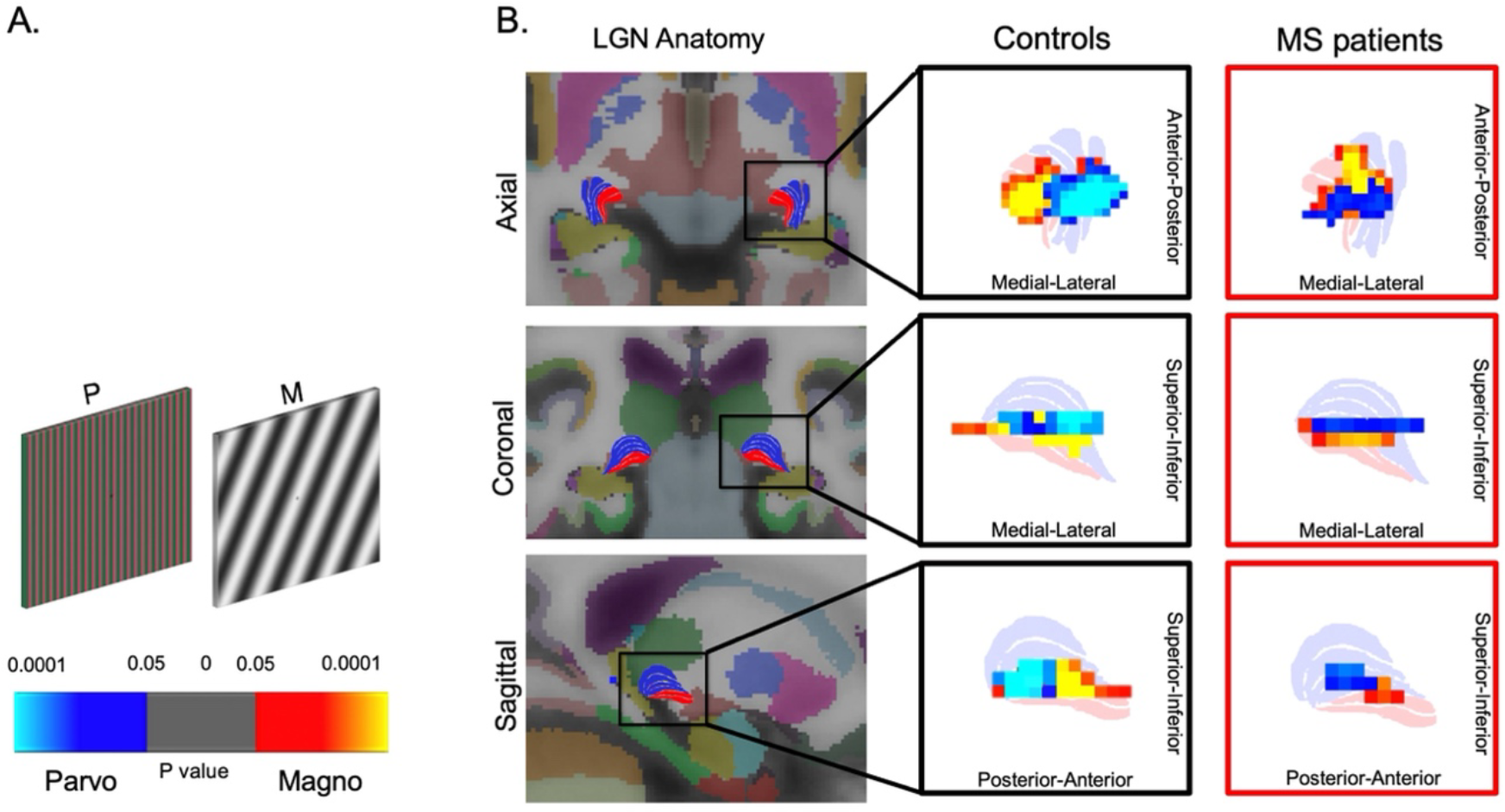
Functional topography of the M and P layers of the LGN in control vs. MS groups. The group functional statistic map identified by contrasting between fMRI responses to the M and P stimuli (paired t-test across all subjects), overlayed on the anatomical map of the LGN. The color bar indicates the statistically significant P-values in red and blue. Only significant voxels were shown in the functional map (A). The first column from the left illustrates the structure of the LGN and its M and P layers from axial, coronal, and sagittal views as a reference, adopted from (Zhang et al., 2015b), which is overlayed on the structural T1 image in MNI space (the M layers are shown in red, and the P layers in blue color). The enlarged spatial pattern of M and P layers in Control and MS patient groups are shown in the second and third columns, respectively (B).

The yellow-red cluster with stronger response to M stimulus is located relatively more inferior (coronal view) and anterior (sagittal view) than the blue voxels which are responded strongly to the P stimulus. In the current study, the topography of identified M and P clusters is consistent with the spatial position of M and P layers in LGN anatomy obtained from brain autopsy (Zhang et al., 2015b). It was also in agreement with other studies which focused on fMRI mapping of LGN subdivisions using M and P stimulation patterns with similar frequency and color characteristics as ours but focused on Healthy (Denison et al., 2014) and glaucoma (Zhang et al., 2016) groups of subjects using different MRI scanners. Notably, the topography of M and P clusters in MS group is highly consistent with those of controls. The axial view shows less distinct M and P functional patterns than sagittal and coronal views, which may be the result of the weak strength of scanners that we used in data collection. In the current study, we used 1.5T and 3T fMRI scanners with isotropic voxel size of 3 *m*^3^and 1.8 *m*^3^, respectively. 3-Tesla scanner data have acceptable spatial resolution compared to similar studies which used 3 and 7 Tesla scanners; however, the spatial resolution of fMRI data acquired by 1.5T scanner is relatively lower than 3T which prevented us providing layer-specific neural response with distinct spatial maps. Thus, increasing number of subjects and recording data with stronger MRI scanner may help to overcome this limitation.

To summarize, the current study shows the malfunction of the whole LGN region and selective impairments in its M and P layers in MS disease. As visual impairment is one of the first symptoms of MS disease, exploring LGN function may be a new potential to become a new way for early diagnosis of MS disease before severe irreversible brain structural damages. Future studies could focus on early-stage MS patients to investigate whether the dysfunction of LGN subdivisions could occur at the early stage. Alongside, increasing the number of subjects can compensate for the effect of brain anatomical variance in statistical analysis in this study, leading to have more reliable and stronger results. Future studies could also consider adding standard chromatic and achromatic vision tests such as Snellen and Ishihara tests. Such data can help as an index for evaluating the severity of visual impairments in MS patients before collecting the visual task-related fMRI data. So, we could better interpret fMRI results by comparing with initial vision test results. Moreover, we used 1.5T and 3Tesla scanners because of their accessibility in hospitals and clinical environments than higher strength scanners. However, future works in this line of research may employ a stronger MRI scanner (7 Tesla or higher). The ability to access higher spatial resolution can help to observe more differentiated layer-specific functional maps in subcortical pathways. Therefore, we could investigate the selective impairment of parallel visual pathways in MS disease. The accurate provoking characteristics of Konio cells are not fully understood yet. To the aim of distinctly provoking chromatic visual pathways (P and K layers in LGN region) by an effective stimulation pattern, in-depth information about Konio cells characteristics in the human brain is needed. So that future works may be able to modify the K stimulation pattern in a way that minimizes provoking P layers at the same time.

## 5. Conclusions

Much is known about the structural damages of primary visual pathways in MS as a result of inflammation and demyelination of nervous system. To the best of our knowledge, discovering functional abnormality in LGN and its subdivisions based on functional MRI remains unaddressed in MS disease. In this study, we assessed LGN function in a group of MS patients compared to healthy controls using task-related fMRI data. M and P cells are organized in distinct spatial positions, while K cells spreads sparsely in form of thin layers between M and P layers. Although fMRI can provide a map of the neural activities in LGN organization, there is no fMRI study on malfunctions of these pathways in MS patients yet.

We measure function of whole LGN, M, P, and K layers by provoking and receiving the BOLD fMRI signals using selective stimulation patterns, i.e., checkerboard, sinusoidal grating black-white, Green-Red and Yellowish green-Bluish purple blocks. We then performed statistical analysis on the preprocessed fMRI data to compare the LGN function of MS patients with those of healthy controls. Our results showed the significant reduction of BOLD signal in the whole LGN body of MS group. Our results showed reduction of average BOLD signal in MS patients in response to M, and P stimulation patterns, suggesting functional impairment of magnocellular and parvocellular pathways in LGN compared to healthy individuals. However, the K stimulation pattern employed in our study did not distinctly provoke K layers resulted in no significant BOLD signal in either MS or healthy group. We thoroughly discussed the limitations of our work in section 4, which could be addressed by future studies in this line of research.

## Acknowledgment

The authors would like to thank the Iranian National Brain Mapping Lab (INBML) for their support and collaboration in data acquisition under the project number P-96185. This research did not receive any specific grant from funding agencies in the public, commercial, or not-for-profit sectors. The authors declare no competing interests.

## Appendix 1.

**Table 1.** MRI imaging parameters for 1.5 Tesla and 3 Tesla scanners.

| Scanner Type | Subjects | Imaging Protocol | Voxel Size (mm<sup>3</sup>) | Slice Number | Slice Thickness (mm) | Flip Angle $\alpha$ (deg) | TR (ms) | TE (ms) | #Volume Number |
| --- | --- | --- | --- | --- | --- | --- | --- | --- | --- |
| 1.5-Tesla,<br>"SIEMENS-MAGNETOM Avanto syngo MR B19" | C1-C5<br>P1-P5 | T<sub>1</sub> mprage sag p2 iso (Structural) | 1×1×1 | 192 | 1 | 8 | 2200 | 2.8 | 1 |
|  |  | GRE field mapping | 3×3×3 | 21 | 3 | 60 |  | 4.76<br>9.52 | Phase:2<br>Amp.:15 |
|  |  | Ep2d fid basic BOLD (Functional) | 3×3×3 | 21 | 3 | 50 | 2000 | 40 | MPK task: 220 |
| 3-Tesla,<br>"SIEMENS MAGNETOM Prisma" | C6-C21<br>P6-P21 | T<sub>1</sub> mprage sag p2 iso (Structural) | 1×1×1 | 176 | 1 | 7 | 1810 | 3.47 | 1 |
|  |  | GRE field mapping | 3×3×3 | 40 | 3 | 60 |  | 5.19<br>7.65 | Phase:2<br>Amp.:15 |
|  |  | Ep2d fid basic BOLD (Functional) | 1.8×1.8×1.8 | 22 | 1.8 | 75 | 2500 | 40 | MPK task: 260<br>LGN loc.: 119 |

## References

Ahmadi, K., Pouretemad, H. R., Esfandiari, J., Yoonessi, A., & Yoonessi, A. (2015). Psychophysical evidence for impaired Magno, Parvo, and Konio-cellular pathways in dyslexic children. Journal of Ophthalmic & Vision Research, 10(4), 433.

Anderson, E. J., Dakin, S. C., & Rees, G. (2009). Monocular signals in human lateral geniculate nucleus reflect the Craik–Cornsweet–O’Brien effect. Journal of Vision, 9(12), 14.

Barnett, M., Graham, E. C., Klistorner, A., Wang, C., Yiannikas, K., You, Y., Garrick, R., Graham, S. L., & Parratt, J. (2017). Progression of retinal ganglion cell loss in multiple sclerosis is associated with new lesions in the optic radiations.

Briggs, F., & Usrey, W. M., (2008). Emerging views of corticothalamic function. Current Opinion in Neurobiology, 18(4), 403–407.

Chen, W., & Zhu, X. (2001). Correlation of activation sizes between lateral geniculate nucleus and primary visual cortex in humans. Magnetic Resonance in Medicine: An Official Journal of the International Society for Magnetic Resonance in Medicine, 45(2), 202–205.

Chen, W., Zhu, X.-H., Thulborn, K. R., & Ugurbil, K. (1999). Retinotopic mapping of lateral geniculate nucleus in humans using functional magnetic resonance imaging. Proceedings of the National Academy of Sciences, 96(5), 2430–2434.

Dacey, D. M. (1999). Primate retina: cell types, circuits and color opponency. Progress in Retinal and Eye Research, 18(6), 737–763.

Denison, R. N., Vu, A. T., Yacoub, E., Feinberg, D. A., & Silver, M. A. (2014). Functional mapping of the magnocellular and parvocellular subdivisions of human LGN. Neuroimage, 102, 358–369.

Dorph-Petersen, K.-A., Caric, D., Saghafi, R., Zhang, W., Sampson, A. R., & Lewis, D. A. (2009). Volume and neuron number of the lateral geniculate nucleus in schizophrenia and mood disorders. Acta Neuropathologica, 117(4), 369.

D’Souza, D. v, Auer, T., Strasburger, H., Frahm, J., & Lee, B. B. (2011). Temporal frequency and chromatic processing in humans: an fMRI study of the cortical visual areas. Journal of Vision, 11(8), 8.

Evangelou, N., Konz, D., Esiri, M. M., Smith, S., Palace, J., & Matthews, P. M. (2001). Size-selective neuronal changes in the anterior optic pathways suggest a differential susceptibility to injury in multiple sclerosis. Brain, 124(9), 1813–1820.

Faro, S. H., Mohamed, F. B., Tracy, J. I., Elfont, R. M., Pinus, A. B., Lublin, F. D., Koenigsberg, R. A., Chen, C. Y., & Tsai, F. Y. (2002). Quantitative functional MR imaging of the visual cortex at 1.5 T as a function of luminance contrast in healthy volunteers and patients with multiple sclerosis. American Journal of Neuroradiology, 23(1), 59–65.

Gareau, P. J., Gati, J. S., Menon, R. S., Lee, D., Rice, G., Mitchell, J. R., Mandelfino, P., & Karlik, S. J. (1999). Reduced visual evoked responses in multiple sclerosis patients with optic neuritis: comparison of functional magnetic resonance imaging and visual evoked potentials. Multiple Sclerosis Journal, 5(3), 161–164.

Haynes, J.-D., Deichmann, R., & Rees, G. (2005). Eye-specific effects of binocular rivalry in the human lateral geniculate nucleus. Nature, 438(7067), 496–499.

Kale, N. (2016). Optic neuritis as an early sign of multiple sclerosis. In Eye and Brain (Vol. 8, pp. 195–202). Dove Medical Press Ltd. https://doi.org/10.2147/EB.S54131

Kastner, S., O’Connor, D. H., Fukui, M. M., Fehd, H. M., Herwig, U., & Pinsk, M. A. (2004). Functional imaging of the human lateral geniculate nucleus and pulvinar. Journal of Neurophysiology, 91(1), 438–448.

Kastner, S., Schneider, K. A., & Wunderlich, K. (2006). Beyond a relay nucleus: neuroimaging views on the human LGN. Progress in Brain Research, 155, 125–143.

Korsholm, K., Madsen, K. H., Frederiksen, J. L., Skimminge, A., & Lund, T. E. (2007). Recovery from optic neuritis: an ROI-based analysis of LGN and visual cortical areas. Brain, 130(5), 1244–1253.

Langkilde, A. R., Frederiksen, J. L., Rostrup, E., & Larsson, H. B. W. (2002). Functional MRI of the visual cortex and visual testing in patients with previous optic neuritis. European Journal of Neurology, 9(3), 277–286.

Martin, P. R., White, A. J. R., Goodchild, A. K., Wilder, H. D., & Sefton, A. E. (1997). Evidence that blue-on cells are part of the third geniculocortical pathway in primates. European Journal of Neuroscience, 9(7), 1536–1541.

McKeefry, D. J., Murray, I. J., & Kulikowski, J. J. (2001). Red–green and blue–yellow mechanisms are matched in sensitivity for temporal and spatial modulation. Vision Research, 41(2), 245–255.

Mullen, K. T., Dumoulin, S. O., & Hess, R. F. (2008). Color responses of the human lateral geniculate nucleus: selective amplification of S-cone signals between the lateral geniculate nucleno and primary visual cortex measured with high-field fMRI. European Journal of Neuroscience, 28(9), 1911–1923.

Mullen, K. T., & Sankeralli, M. J. (1999). Evidence for the stochastic independence of the blue-yellow, red-green and luminance detection mechanisms revealed by subthreshold summation. Vision Research, 39(4), 733–745.

Mullen, K. T., Thompson, B., & Hess, R. F. (2010). Responses of the human visual cortex and LGN to achromatic and chromatic temporal modulations: an fMRI study. Journal of Vision, 10(13), 13.

Murav’eva, S. v, Deshkovich, A. A., & Shelepin, Y. E. (2009). The human magno and parvo systems and selective impairments of their functions. Neuroscience and Behavioral Physiology, 39(6), 535–543.

O’Connor, D. H., Fukui, M. M., Pinsk, M. A., & Kastner, S. (2002). Attention modulates responses in the human lateral geniculate nucleus. Nature Neuroscience, 5(11), 1203–1209.

Pawlitzki, M., Horbrügger, M., Loewe, K., Kaufmann, J., Opfer, R., Wagner, M., Al-Nosairy, K. O., Meuth, S. G., Hoffmann, M. B., & Schippling, S. (2020). MS optic neuritis-induced long-term structural changes within the visual pathway. Neurology-Neuroimmunology Neuroinflammation, 7(2).

Pihl-Jensen, G., Schmidt, M. F., & Frederiksen, J. L. (2017). Multifocal visual evoked potentials in optic neuritis and multiple sclerosis: a review. Clinical Neurophysiology, 128(7), 1234–1245.

Planche, V., Su, J. H., Mournet, S., Saranathan, M., Dousset, V., Han, M., Rutt, B. K., & Tourdias, T. (2020). White-matter-nulled MPRAGE at 7T reveals thalamic lesions and atrophy of specific thalamic nuclei in multiple sclerosis. Multiple Sclerosis Journal, 26(8), 987–992.

Rocca, M. A., & Filippi, M. (2007). Functional MRI in multiple sclerosis. Journal of Neuroimaging, 17, 36S–41S.

Rombouts, S., Lazeron, R. H. C., Scheltens, P., Uitdehaag, B. M. J., Sprenger, M., Valk, J., & Barkhof, F. (1998). Visual activation patterns in patients with optic neuritis: An f MRI pilot study. Neurology, 50(6), 1896–1899.

Russ, M. O., Cleff, U., Lanfermann, H., Schalnus, R., Enzensberger, W., & Kleinschmidt, A. (2002). Functional magnetic resonance imaging in acute unilateral optic neuritis. Journal of Neuroimaging, 12(4), 339–350.

Schmielau, F., & Singer, W. (1977). The role of visual cortex for binocular interactions in the cat lateral geniculate nucleus. Brain Research.

Schneider, K. A., Richter, M. C., & Kastner, S. (2004). Retinotopic organization and functional subdivisions of the human lateral geniculate nucleus: a high-resolution functional magnetic resonance imaging study. Journal of Neuroscience, 24(41), 8975–8985.

Sharma, S., Chitranshi, N., vander Wall, R., Basavarajappa, D., Gupta, V., Mirzaei, M., Franzco, S. L. G., Klistorner, A., & You, Y. (2021). Trans-synaptic degeneration in the visual pathway: neural connectivity, pathophysiology, and clinical implications in neurodegenerative disorders. Survey of Ophthalmology.

Sherman, S. M., & Guillery, R. W. (2002). The role of the thalamus in the flow of information to the cortex. Philosophical Transactions of the Royal Society of London. Series B: Biological Sciences, 357(1428), 1695–1708.

Sun, Y., Huang, W., Li, F., Li, H., Wang, L., Huang, Y., & Zhang, X. (2019). Subcortical visual pathway may be a new way for early diagnosis of glaucoma. Medical Hypotheses, 123, 47–49.

Talman, L. S., Bisker, E. R., Sackel, D. J., Long Jr, D. A., Galetta, K. M., Ratchford, J. N., Lile, D. J., Farrell, S. K., Loguidice, M. J., & Remington, G. (2010). Longitudinal study of vision and retinal nerve fiber layer thickness in multiple sclerosis. Annals of Neurology, 67(6), 749–760.

Tian, D.-C., Su, L., Fan, M., Yang, J., Zhang, R., Wen, P., Han, Y., Yu, C., Zhang, C., & Ren, H. (2018). Bidirectional degeneration in the visual pathway in neuromyelitis optica spectrum disorder (NMOSD). Multiple Sclerosis Journal, 24(12), 1585–1593.

Toosy, A. T., Werring, D. J., Bullmore, E. T., Plant, G. T., Barker, G. J., Miller, D. H., & Thompson, A. J. (2002). Functional magnetic resonance imaging of the cortical response to photic stimulation in humans following optic neuritis recovery. Neuroscience Letters, 330(3), 255–259.

Werring, D. J., Bullmore, E. T., Toosy, A. T., Miller, D. H., Barker, G. J., MacManus, D. G., Brammer, M. J., Giampietro, V. P., Brusa, A., & Brex, P. A. (2000). Recovery from optic neuritis is associated with a change in the distribution of cerebral response to visual stimulation: a functional magnetic resonance imaging study. Journal of Neurology, Neurosurgery & Psychiatry, 68(4), 441–449.

Wesner, M. F., & Brazeau, J. (2019). The Psychophysical Assessment of Hierarchical Magno-, Parvo-and Konio-Cellular Visual Stream Dysregulations in Migraineurs. Eye and Brain, 11, 49.

Yoonessi, A., & Yoonessi, A. (2011). Functional assessment of magno, parvo and konio-cellular pathways; current state and future clinical applications. Journal of Ophthalmic & Vision Research, 6(2), 119.

Zhang, P., Wen, W., Sun, X., & He, S. (2016). Selective reduction of fMRI responses to transient achromatic stimuli in the magnocellular layers of the LGN and the superficial layer of the SC of early glaucoma patients. Human Brain Mapping, 37(2), 558–569.

Zhang, P., Zhou, H., Wen, W., & He, S. (2015a). Layer-specific response properties of the human lateral geniculate nucleus and superior colliculus. Neuroimage, 111, 159–166.

Zhang, P., Zhou, H., Wen, W., & He, S. (2015b). Layer-specific response properties of the human lateral geniculate nucleus and superior colliculus. Neuroimage, 111, 159–166.

Zivadinov, R., Bergsland, N., Dolezal, O., Hussein, S., Seidl, Z., Dwyer, M. G., Vaneckova, M., Krasensky, J., Potts, J. A., & Kalincik, T. (2013). Evolution of cortical and thalamus atrophy and disability progression in early relapsing-remitting MS during 5 years. American Journal of Neuroradiology, 34(10), 1931–1939.

